# Insights into cisplatin-induced neurotoxicity and mitochondrial dysfunction in *Caenorhabditis elegans*

**DOI:** 10.1101/2021.06.03.445826

**Authors:** Carmen Martínez-Fernández, Milana Bergamino, David Brena, Natascia Ventura, Sebastian Honnen, Alberto Villanueva, Ernest Nadal, Julián Cerón

**Affiliations:** Genes, Diseases, and Therapies Program, Bellvitge Biomedical Research Institute (IDIBELL), L’Hospitalet de Llobregat, 08908 Barcelona, Spain; Department of Medical Oncology, Breast Cancer Unit, Catalan Institute of Oncology, Hospital Duran i Reynals, Avda Gran via, 199-203, L’Hospitalet, 08908, Barcelona, Spain; IUF-Leibniz Research Institute for Environmental Medicine, 40225 Düsseldorf, Germany; Institute of Toxicology, Medical Faculty, Heinrich Heine University, Universitätsstraße 1, D-40225 Düsseldorf, Germany; Group of Chemoresistance and Predictive Factors, Subprogram Against Cancer Therapeutic Resistance, Catalan Institute of Oncology, Oncobell Program, Bellvitge Biomedical Research Institute (IDIBELL), L’Hospitalet del Llobregat, Barcelona, Spain; Medical Oncology Department, Catalan Institute of Oncology, L’Hospitalet, Barcelona Spain

## Abstract

Cisplatin is the most common drug in first-line chemotherapy against solid tumors. We and others have previously used the nematode *Caenorhabditis elegans* to identify genetic factors influencing the sensitivity and resistance to cisplatin. In this study, we take advantage of *C. elegans* to explore cisplatin effects on mitochondrial functions and investigate cisplatin-induced neurotoxicity through a high-resolution semi-automated system for evaluating locomotion. Firstly, we report that a high-glucose diet sensitizes *C. elegans* to cisplatin at the physiological level and that mitochondrial CED-13 protects the cell from cisplatin-induced oxidative stress. Additionally, by assessing mitochondrial function with a Seahorse Analyzer, we observed a detrimental additive effect of cisplatin and glucose in mitochondrial respiration. Secondly, since we previously found that catechol-*O*-methyltransferases (involved in dopamine degradation) were upregulated upon cisplatin exposure, we studied the protective role of the FDA-approved drug dopamine against cisplatin-induced neurotoxicity. To implement the use of the Tierpsy Tracker system for measuring neurotoxicity in *C. elegans*, we showed that abnormal displacements and body postures in *cat-2* mutants, which have the dopamine synthesis pathway disrupted, can be rescued by adding dopamine. Then, we used such a system to demonstrate that dopamine treatment protects from the dose-dependent neurotoxicity caused by cisplatin.

## INTRODUCTION

Cis-Diammine Dichloride Platinum (CDDP), also known as cisplatin, is one of the most used platinum derivatives and the one with the highest therapeutic efficacy in a series of solid tumors, including testicular, ovarian, cervical, bladder, head and neck, and lung cancer (Amable, 2016; Dasari and Bernard Tchounwou, 2014). The most remarkable example is testicular cancer, in which cisplatin provides the cure for more than 80% of the patients (Gonzalez-Exposito et al., 2016). Unfortunately, despite its effectiveness, oncologists need to deal with three main inconveniences associated with its use which ultimately results in therapy failure and increased mortality: *(i)* the tumor-acquired resistance (Amable, 2016), *(ii)* the intrinsic resistance of many patients and *(iii)* the toxic side-effects (Kelland, 2007). Therefore, identification of factors predicting dosage sensitivity to therapy, and the finding of novel resensitizing therapeutic approaches are essential for effective treatments with cisplatin.

Besides the genetic background, the metabolic profile also influences cancer progression and outcome, and response to chemotherapy. Metabolic disorders, such as Type 2 Diabetes Mellitus (T2DM), negatively affect the clinical outcome of cancer patients (Barone et al., 2008; Harding et al., 2015; Tsilidis et al., 2015). Recently, it has been suggested that fasting plasma glucose (FPG) is a predictor of survival in non-small cell lung cancer (NSCLC) patients treated with concurrent chemoradiotherapy since higher FPG correlates with a lower overall survival rate (Bergamino et al., 2019). Therefore, although a good metabolic control would improve cancer outcome, there are conflicting data providing the efficiency of antidiabetic compounds on cancer mortality (Lin et al., 2015; Ranc et al., 2014; Shlomai et al., 2016).

The unspecific mode of action of cisplatin works as a double-edged sword affecting both tumoral and normal cells. CDDP undergoes aquation when entering the cell, becoming more reactive to interact with a broad range of cellular targets, both in the nucleus and cytoplasm (Kelland, 2007). Among the cytotoxic consequences of cisplatin, the formation of DNA-cisplatin adducts and the production of reactive oxygen species, which leads to homeostasis imbalance and ultimately provoke apoptosis, are considered the canonical mechanisms of action (Wang and Lippard, 2005). Thus, the need to reduce toxic side effects implies a dose-limiting outcome in cisplatin-based therapies.

The most common side effect is ototoxicity, affecting more than 60% of pediatric cancer patients, peripheral nervous system toxicity, and nephrotoxicity. This particular toxicity leads to partial or even complete hearing loss, compromising the language and cognitive development of these patients (Ross et al., 2009). Interestingly, Catechol-*O*-methyltransferase (COMT), acylphosphatase 2 (ACYP2), and thiopurine methyltransferase (TPMT) genetic variants have been related to ototoxicity (Ross et al., 2009). However, further reports show contradictory data (Thiesen et al., 2017) and the mechanisms undergoing this common pathophysiology remain still obscure.

*Caenorhabditis elegans* has been successfully used as an experimental organism since the 1970s, providing several advantages over other models. This nematode is a well-established system to investigate neuromodulatory pathways *in vivo* (Van Damme et al., 2021) and the effect of genotoxic drugs in a pluricellular context (Honnen, 2017; Kaletta and Hengartner, 2006). Particularly, we and others have recently demonstrated the value of *C. elegans* to explore genetic, cellular, and molecular factors involved in cisplatin response (Hemmingsson et al., 2010; García-Rodríguez et al., 2018; Wellenberg et al., 2021). Here, exploiting our previous methodologies, along with CRISPR-Cas9 technology and semi-automated platforms (Seahorse XFe96 Analyzer and Tierpsy Tracker), we further investigated the effects of cisplatin-based therapies. On the one hand, we observed that a glucose-enriched diet negatively impacts cisplatin response, particularly affecting mitochondrial function. Moreover, we demonstrate the protective role of the BH3-only protein CED-13 against cisplatin-induced mtROS. On the other hand, we found that dopamine protects from cisplatin-induced neurotoxicity on animal locomotion.

## RESULTS

### High-glucose diet influences the animal response to cisplatin

In a previous study, we described a dose-response effect of cisplatin on *C. elegans* body length during postembryonic development (García-Rodríguez et al., 2018). In the same report, we pointed out the insulin/IGF-1 signaling (IIS) pathway as a critical regulator in the animal response to cisplatin, suggesting the relevance of metabolism in cisplatin chemosensitivity. To investigate whether hyperglycemic conditions impact cisplatin response in the nematode, we assessed the effect of a high-glucose diet on animals’ body length during larval development under our standard cisplatin conditions (60 μg/mL). We performed a dose-response assay including glucose concentrations in *C. elegans* whole-body extract of 10-15 mM, resembling the glucose levels of diabetic patients under poor glucose control (Schlotterer et al., 2009). Our data showed that high-glucose diet does not cause a major effect on *C. elegans* development, but when co-administered with cisplatin, high glucose concentrations (40 and 80 mM) sensitize the animals to cisplatin (**Fig. 1**). Interestingly, low glucose supplementation (10 mM) slightly protects the animal from the drug effects.

**Figure 1:**
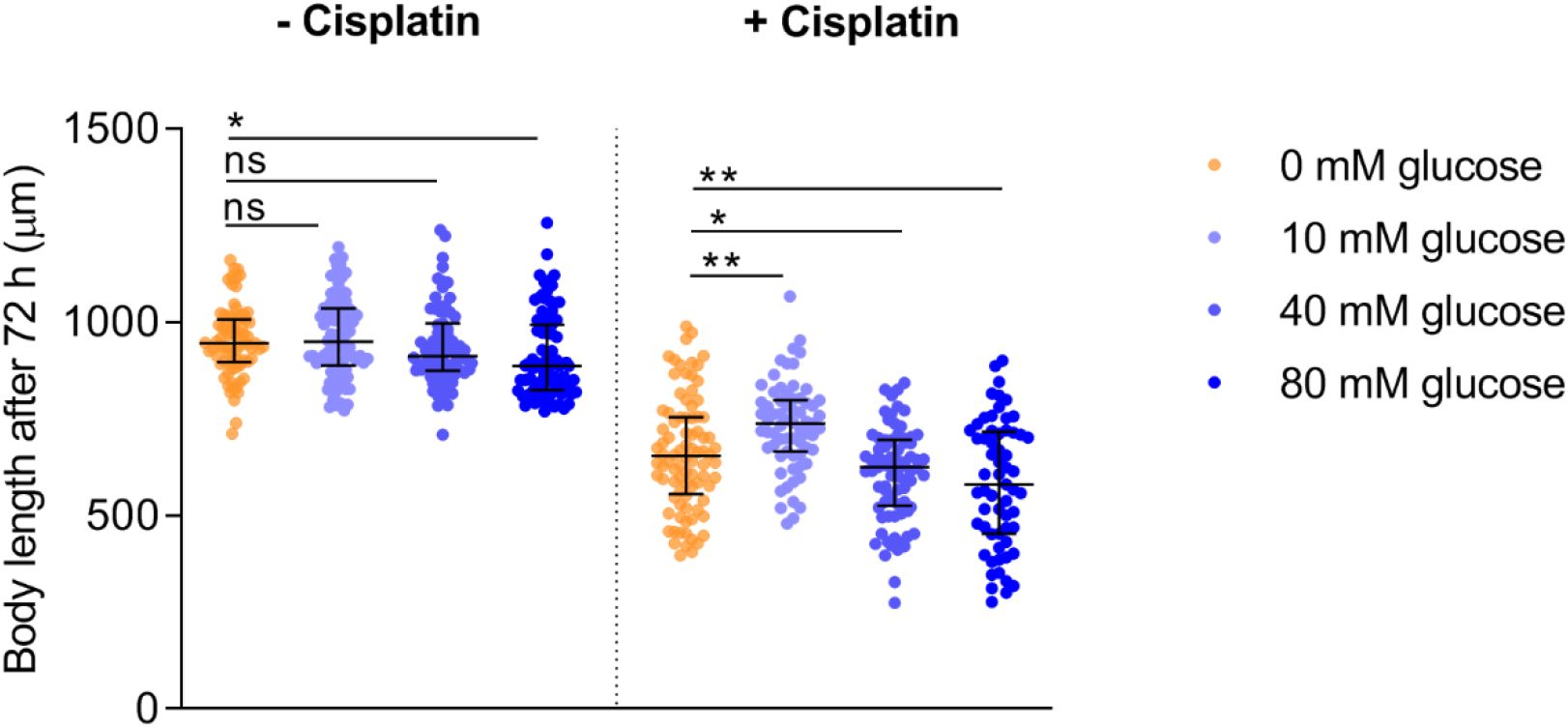
Glucose supplementation enhances the cisplatin effect on *C. elegans* body length. The graph shows animals’ body length 72-hours post seeding L1 animals at 20°C, fed with different glucose concentrations, and exposed or not to 60 μg/mL of cisplatin. Bars indicate the median and interquartile range and dots body length values of individual animals. Two independent experiments were performed, and each condition was measured in duplicates, analyzing 50 animals per condition. *, ** and ns mean p<0.1, p<0.01, and no significant, respectively. Statistics were analyzed by ordinary one-way ANOVA (Holm-Sidak’s test).

### The combination of mtROS and cisplatin causes an additive adverse effect

High-glucose levels alter mitochondrial properties and functions in *C. elegans* (Alcántar-Fernández et al, 2019). We investigated if the observed synergistic effect of glucose and cisplatin in reducing the body length could occur through oxidative stress induction and ROS generation at the mitochondrial level. Thus, we tested if cisplatin displays a cumulative effect with the oxidant paraquat, a potent mitochondrial ROS (mtROS) generator. First, we assessed the dose-dependent impact of paraquat (PQ) on *C. elegans* development (**Fig. 2A**). Thus, a mild toxic PQ dose (0.1 mM), high enough to provoke a significant reduction in body length but allowing the detection of cisplatin-induced modulatory effect was selected for subsequent experiments. Finally, we noticed that cisplatin (60 μg/ml) together with paraquat (0.1 mM) have a increased impact on body length (**Fig. 2B**) similar to that observed with high-glucose diet (**Fig. 1**).

**Figure 2:**
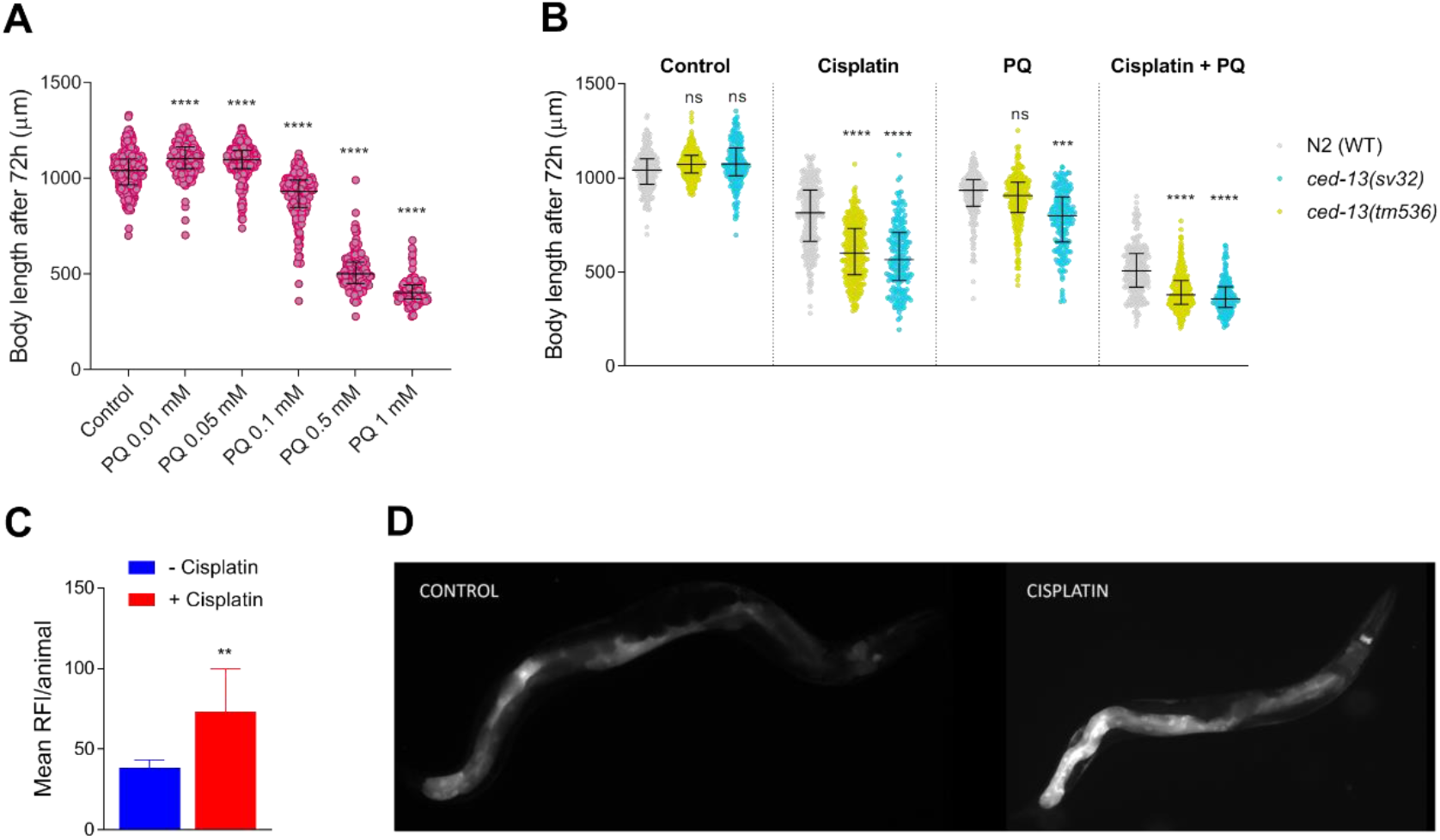
Cisplatin induces mtROS and activates mitochondrial damage response pathways. **A.** Dose-response curve showing paraquat effect on wild-type animals’ body-length. **B.** Additive effect of cisplatin and PQ in WT and *ced-13* mutants (*sv32* and *tm536*). Statistics were analyzed by one-way ANOVA (Krustal-Wallis and Dunn’s tests). ***, **** and ns mean, *p*<0.001, *p*<0.0001 and non-significant, respectively, compared to wild-type in the same drug condition. Three independent experiments were performed. **C.** *hsp-6p::GFP* relative fluorescence intensities (RFI) are represented for in control and cisplatin-treated animals. Bars represent the mean of two independent experiments (twenty animals per experiment) and lines the standard deviation. Statistics were performed using Student’s t-test, ** means p<0.01. **D.** Representative images of animals expressing *hsp*-*6p::GFP* under control and cisplatin conditions.

The *C. elegans* BH3-only protein CED-13 promotes cell survival in response to mtROS, instead of inducing apoptosis (Yee et al., 2014). Moreover, as we have previously described, CED-13 protects *C. elegans* from cisplatin-induced toxicity (García-Rodríguez et al., 2018) (**Fig. 2B**). To confirm that cisplatin and paraquat synthetic toxicity occur through the production of mtROS, we used two different *ced-13* mutant alleles, *sv32* and *tm536*, harboring 1304-bp and 523-bp deletions, respectively. Interestingly, we observed that CED-13 also protects against cisplatin and paraquat synergistic toxic effect (**Fig. 2B**). These results suggest that toxicity caused by cisplatin in combination with glucose or PQ occurs by oxidative stress induction affecting the mitochondria.

To further corroborate the adverse effect of cisplatin on mitochondrial functions, we evaluated the activation of the mitochondrial unfolded protein response (UPR^mt^) pathway. The UPR^mt^ pathway is induced in response to misfolded or unassembled proteins within the mitochondria or when mitochondrial respiratory complexes are imbalanced. To assess the UPR^mt^ activation, we quantified the expression of HSP-6, a chaperone commonly used to rate the UPR^mt^ pathway activation (Jovaisaite et al., 2014). Animals treated with our standard concentration of cisplatin increased the expression of the *hsp*-*6p::GFP* reporter (**Fig. 2D**). This observation supports the hypothesis that cisplatin may have a detrimental effect on mitochondrial functions.

### Impact of cisplatin and glucose in mitochondrial respiration

The disruption of the stoichiometric balance between components of mitochondrial respiratory complexes (OXPHOS complexes I, III, IV and V) is one of the signals triggering UPR^mt^ (Jovaisaite et al., 2014). To investigate the impact of cisplatin and high-glucose levels on the activity of OXPHOS complexes, we assessed the mitochondrial respiration of L3 animals using the Seahorse XFe96 respirometer, including paraquat-treated animals as a control group. The Seahorse XFe96 facilitates the measurement of the oxygen consumption rate (OCR), a measure of mitochondrial function and energy production rate, before and after the sequential injection of two mitochondrial complex inhibitors: carbonyl cyanide-4-(trifluoromethoxy) phenylhydrazone (FCCP) and sodium azide (Koopman et al., 2001) Thus, allowing to extract the basal respiration, the maximal respiratory capacity, and the spare respiratory capacity values (**Fig. 3A**). We measured the OCR in L3 larvae after 24-hours exposure to the correspondent treatment. As expected, cisplatin dramatically affected the basal respiration (**Fig. 3B, C**), the maximal respiratory capacity (**Fig. 3B, D**). and the spare capacity (**Fig. 3E**). Interestingly, we observed an additive effect of cisplatin with mtROS producers (glucose and PQ) on the maximal respiratory capacity and the spare capacity, highlighting their additive detrimental effects on mitochondrial respiration.

**Figure 3:**
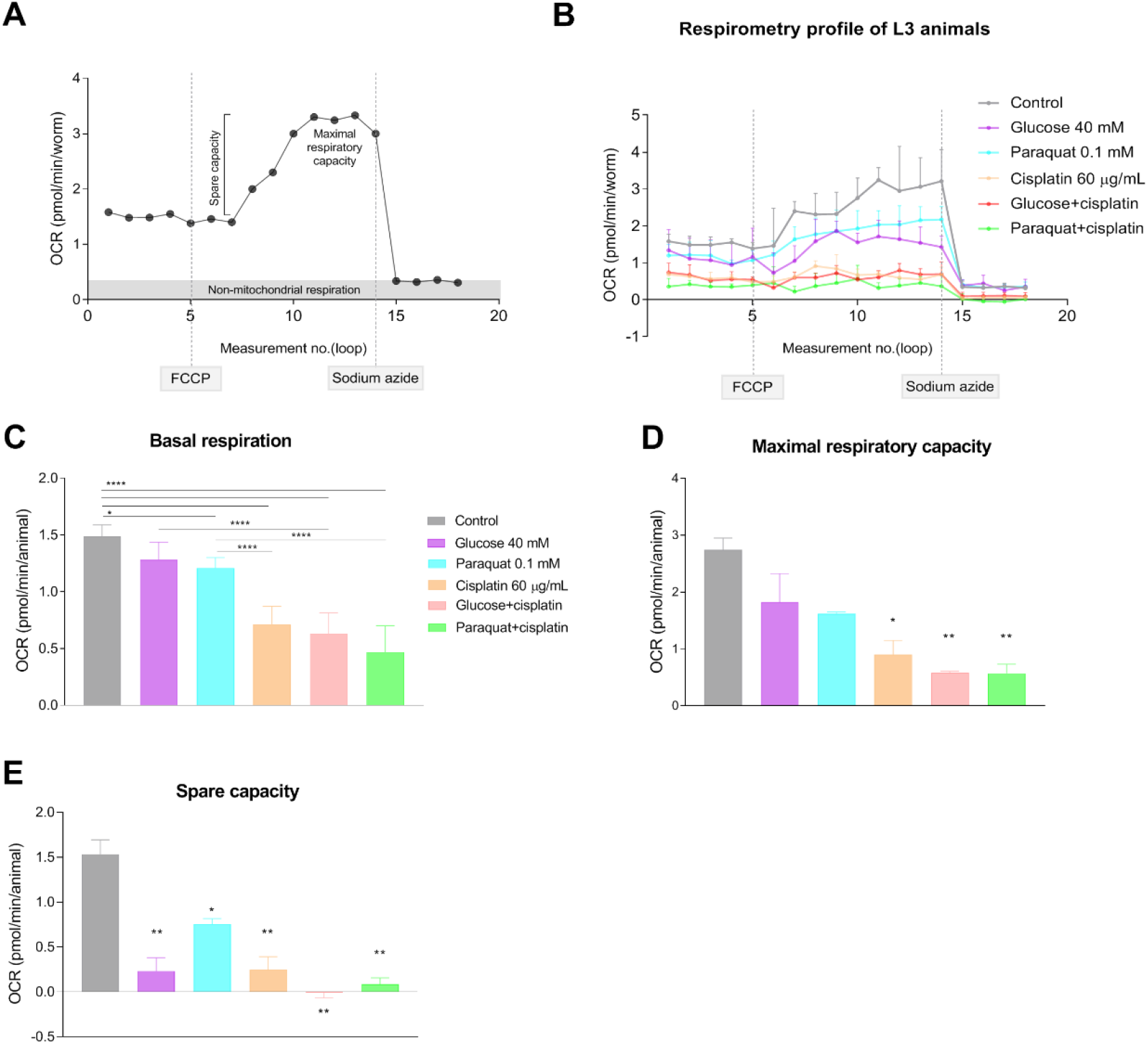
*C. elegans* respirometry evaluation. **A.** Typical OCR *C. elegans* respirometry profile in adult animals. Modified from (Koopman et al., 2016). Before drug injections, the respirometer informs about *basal respiration*. Then, FCCP injection disrupts the mitochondrial membrane potential and ATP synthesis while still allowing proton pumping, electron transporting, and oxygen consumption. Thus, FCCP enables the measurement of *maximal respiratory capacity*. The extraction of the *basal respiration* from the *maximal respiratory capacity* results in the *spare respiratory capacity*, a value indicating the organism ability to respond to increasing energy demands. Finally, injection of sodium azide blocks both cytochrome c oxidase (complex IV) and the ATP synthase (complex V), thereby shutting down the whole electron transport chain and allowing to distinguish non-mitochondrial oxygen consuming processes. **B.** OCR profile of *C. elegans* L3 stage larvae in control and treated conditions. Connected points represent the median of the measures of each condition at a given loop and lines represent standard deviation. Discontinuous lines indicate FCCP and sodium azide injections, respectively. Median and interquartile range are represented by bars and error bars, respectively, for basal respiration (**C**), maximal respiratory capacity (**D**) and spare capacity (**E**). One-way ANOVA (Holm-Sidak’s and Dunn’s tests) were used to compare statistical differences between groups. Experiment was performed twice. **** p<0.0001, ** p<0.01, * p<0.1. Data were analyzed using Agilent Seahorse XFe96 Analyzer, Seahorse Wave Desktop software and GraphPad Prism 8.0.

### COMT genes are involved in dopamine-dependent processes

One of the intriguing genes that showed upregulation upon cisplatin exposure in our previous publication was *comt-4*, which encodes a catechol-*O*-methyltransferase related to the degradation of catecholamines, including dopamine (DA) (García-Rodríguez et al., 2018). Since genetic variants of the human COMT gene have been linked to ototoxicity in cisplatin-treated children (Ross et al., 2009), we decided to investigate in our model the role of COMT genes in cisplatin-induced neurotoxicity.

*C. elegans* COMT family presents five members (*comt*-*1* to *comt*-*5*), all containing the catechol-*O*-methyltransferase domain. Human COMT gene encodes for two similar isoforms, S-COMT and M-COMT, with 100% sequence similarity but with different subcellular localization. BLAST alignment using S-COMT as reference sequence shows high sequence similarity with *C. elegans* COMT members, particularly at the N-terminal end (**Fig. S1**). Interestingly, most of the binding sites for S-COMT ligands are conserved or partially conserved along all the *C. elegans* COMTs, suggesting the conservation of functional roles. COMT-4 possesses the highest percentage of aminoacid identity per query cover (**Table S1**) with human S-COMT. A *comt*-*4* endogenous transcriptional reporter generated by Nested CRISPR (Vicencio et al., 2019) display expression in neuronal cells as expected from its functional role and the transcriptomic data from GExplore (Cao et al., 2017; Hutter and Suh, 2016).

Since the five *C. elegans* COMT family members could present certain functional redundancy, besides *comt*-*4* we also investigated *comt*-*3* and *comt*-*5*, because of their higher expression at postembryonic stages compared to *comt*-*1* and *comt*-*2* (Hutter and Suh, 2016). Thus, we generated deletion alleles for the three genes by CRISPR-Cas9 (*comt*-*3(cer130)*, *comt*-*4(cer126)* and *comt*-*5(cer128)*) (**Fig. S2**).

First, we examined body length, which is negatively regulated by dopamine (Nagashima et al., 2016), in the three COMT mutants. As experimental control, we used a null allele for *cat*-*2*, a gene encoding a tyrosine hydroxylase required for dopamine synthesis. *cat*-*2(n4547)* mutant present reduced levels of dopamine (Smith et al., 2019), thus affecting dopamine-dependent behavioral and morphological effects (Omura et al., 2012; Smith et al., 2019). Body length of synchronized animals were measured at 96 hours post-seeding. As expected, *cat*-*2(n4547)* dopamine-defective animals were larger than wild-type animals. In contrast, *comt*-*5(cer126)* animals, and a triple mutant strain harboring *comt*-*3*, *comt*-*4* and *comt*-*5* deletion alleles, were shorter than wild-type (**Fig. S3**). This result suggests that *comt*-*5* has a greater implication in dopamine catabolism than *comt*-*3* and *comt*-*4*. Still, we cannot discard other impacts of COMT genes and functional redundancies in distinct dopamine-regulated processes.

### COMT mutants show behavioral phenotypes even in the absence of cisplatin

To study cisplatin-induced neurotoxicity in *C. elegans* we evaluated different dopamine-dependent phenotypes in control and cisplatin conditions. However, none of these standard assays clearly showed differences between WT and COMT mutants. Thus, we established a methodology to automatically track animals using the Tierpsy Tracker (Javer et al., 2018a) (**Fig. 4A**). This system combines the throughput of multi-worm tracking with the resolution of single worm movements, allowing the extraction of detailed phenotypic fingerprints from a population (Stephens et al., 2008).

**Figure 4:**
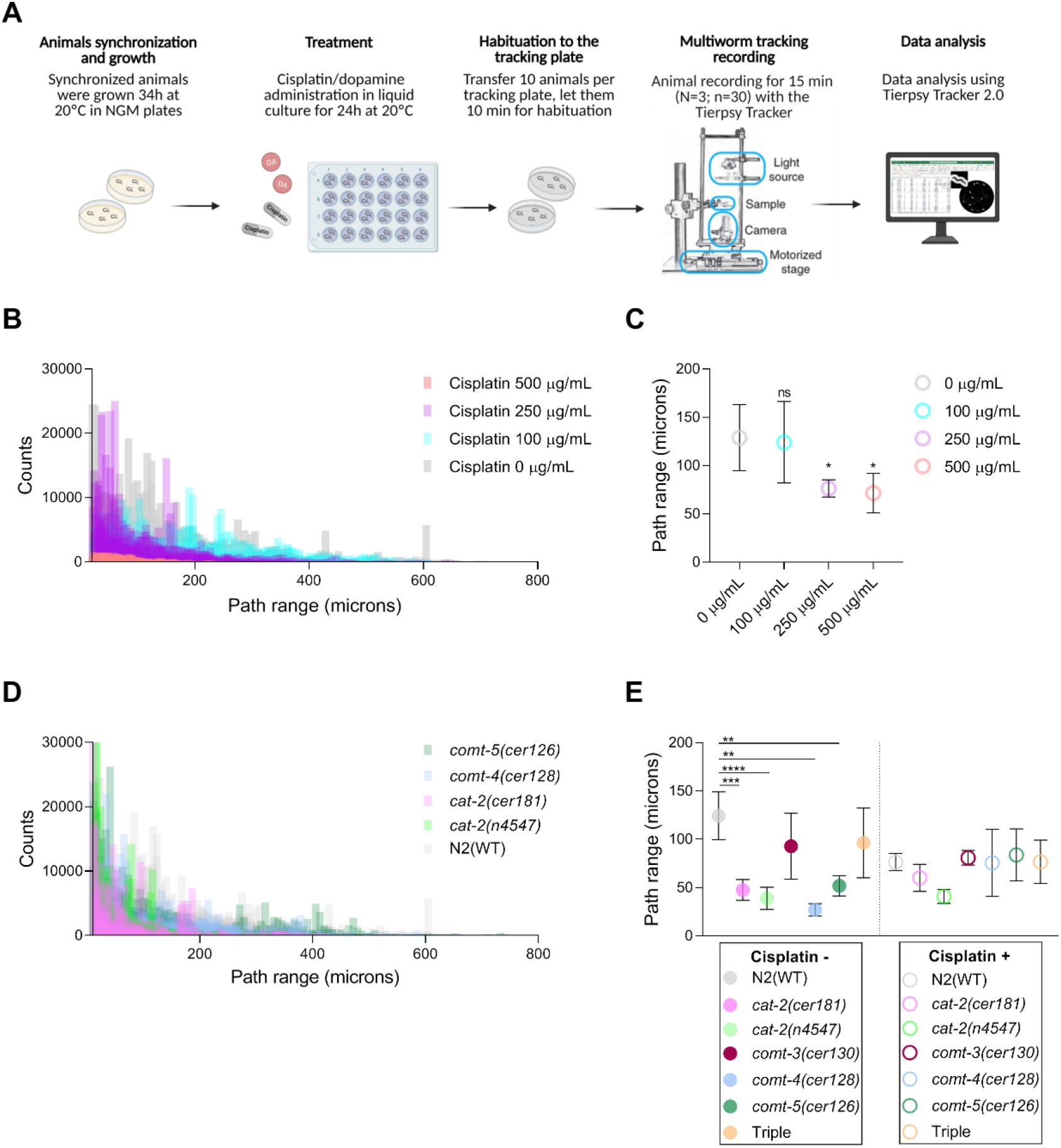
Neurotoxic evaluation using the Tierpsy Tracker in wild type and dopamine signaling-related mutants. **A.** Schematic representation of the experimental flow followed to evaluate neuronal functions under control and treated (cisplatin and dopamine) conditions. Created with Biorender.com. **B.** Histograms show the dose-dependent effect of cisplatin on animal path range in wild type animals. **C.** Circles represent the mean path range of animals exposed to cisplatin; back lines represent standard deviation. **D.** Histograms represent the path range profile of wild type, *cat*-*2*, *comt*-*4* and *comt*-*5* mutants. **E.** Circles represent the mean path range of animals exposed or not to 250 μg/mL cisplatin. Black lines represent standard deviation of two independent experiments. *, **, ***, **** mean p<0.1, p<0.01, p<0.001 and p<0.0001. Statistics was performed by one-way ANOVA (Krustal-Wallis and Dunn’s tests).

We determined cisplatin conditions leading to an altered behavioral phenotype in adult wild-type animals. L4 animals were treated with distinct cisplatin concentrations (100, 250 and 500 μg/mL) for 24 hours. Then, nematodes were transferred to tracking plates and recorded for 15 minutes. Finally, we processed data files using Tierpsy Tracker 2.0 software (Javer et al., 2018b) (**Fig. 4A**). Cisplatin produced a dose-dependent decrease in the overall path range, defined as the distance of the animal’s midbody from the path centroid measured in microns (**Fig. 4B, C and Movie S1**). We used 250 μg/mL as the standard concentration of cisplatin on plates because it was the highest possible dose that didn’t affect viability.

Thus, we used the tracker system to study the impact of cisplatin (250 μg/mL) on the locomotor activity of distinct mutant backgrounds. As a control, we included two *cat*-*2* deletion alleles, *cer181* and *n4547*. By CRISPR-Cas9, we generated the allele *cer181* because the *n4547* allele affects the UTR of an additional locus *(pqn*-*85)*. These *cat*-*2* mutants were expected to show impaired locomotion compared to wild-type animals due to a hampered dopamine signaling (Omura et al., 2012; Smith et al., 2019). We subjected both *cat*-*2* mutants along with COMT mutants (*comt*-*3(cer130)*, *comt*-*4(cer126)*, *comt*-*5(cer128)* and triple mutant) and wild-type animals to path range evaluation in control and cisplatin conditions. Significant reduction in the traveled distances compared to wild-type was evidenced in untreated conditions not only for *cat*-*2* mutants (**Movie S2**) but also for *comt*-*4* and *comt*-*5* deletion alleles (**Fig. 4D, E**). Path ranges were not further altered in any mutant strain under cisplatin exposure (**Fig. 4E**). Thus, although we expected wild-type locomotion in COMT null mutants, they displayed a behavioral phenotype too strong in control conditions to study their sensitivity to cisplatin in our experimental conditions.

### Dopamine protects from cisplatin-induced neurotoxicity

Dopamine-deficient mutants, *cat*-*2(cer181)* and *cat*-*2(n4547)*, traveled shorter distances than animals with normal levels of dopamine (**Fig. 4D, E and Movie S2**). To study the neuroprotective effect of DA in cisplatin-treated animals, we first tested the efficacy of exogenous DA (5 and 10 Mm) in live animals by rescuing the locomotor phenotypes of *cat*-*2* mutants. Low DA supplementation (5 mM) was enough to increase *cat*-*2* mutants path ranges up to wild-type values. Interestingly, we observed a detrimental effect of DA at 10 mM in the WT background (**Fig. 5A**), further supporting that proper DA levels are essential to maintain normal displacements. The Tierpsy Tracker not only reports locomotory features but also the range of shapes adopted by nematodes (Javer et al., 2018b). The posture of the worm can be reconstructed as a summation of eigenworms or eigenprojections. We consider six eigenprojections (*α*1- *α*6) which can almost completely describe the natural worm posture by measuring the worm curvature (**Fig. S4A**). We noticed that 5- and 10-mM exogenous DA could also rescue altered body posture, particularly eigenprojections *α*1, *α*2, and *α*6 to wild-type values (**Fig. S4B, C**).

**Figure 5:**
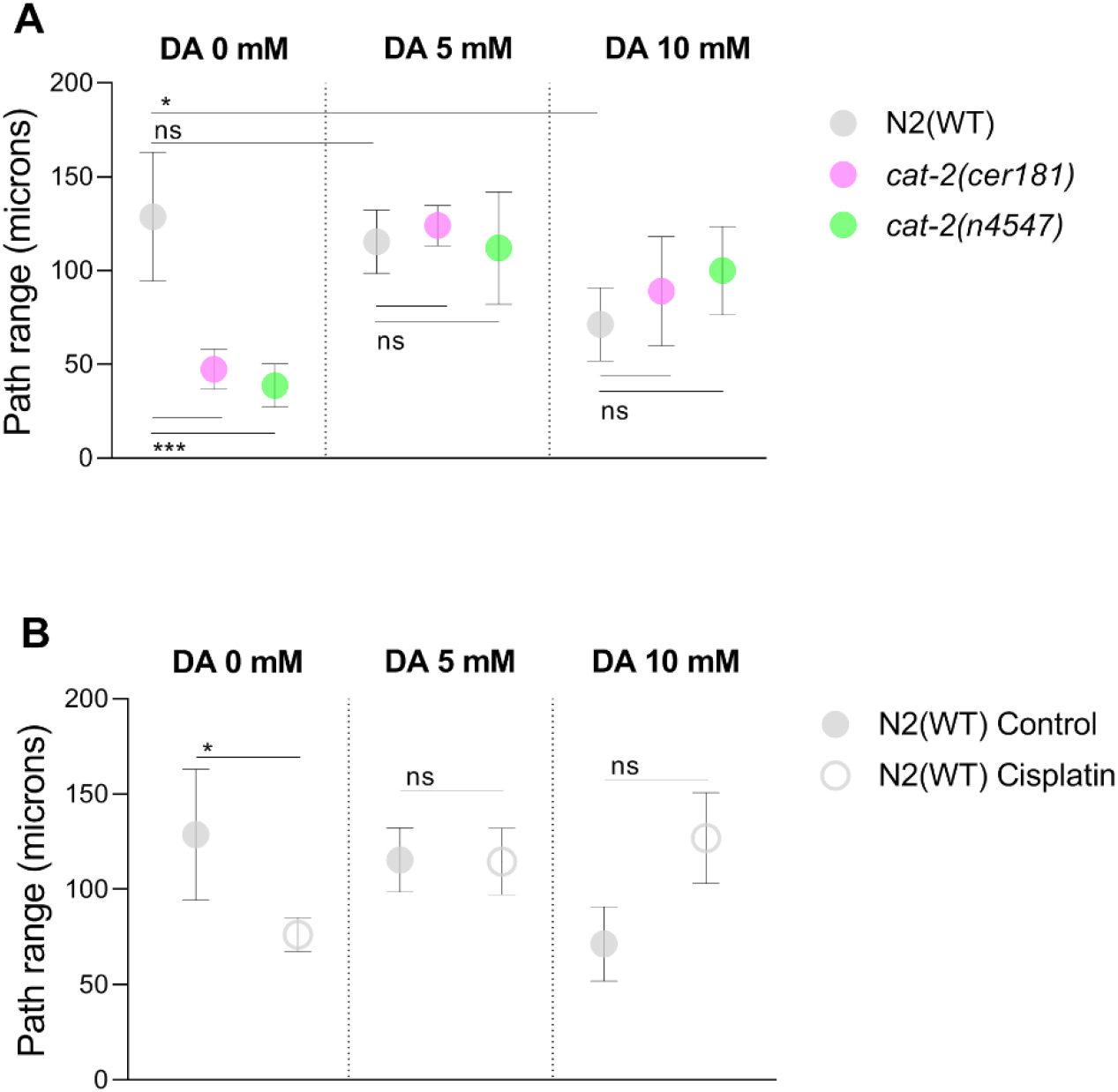
Dopamine influences path range and protect from cisplatin-induced neurotoxicity. Dopamine rescues the behavioral defects of low-dopamine mutants (**A**) and cisplatin-exposed animals (**B**). Circles represent the mean path range of animals control or exposed to cisplatin. Black lines represent standard deviation. DA concentration is indicated at the top of the graph. Two biological replicates were performed. *, ***, **** and ns mean p<0.01, p<0.001, p<0.0001 and not significant, respectively. Statistics was performed by one-way ANOVA (Kruskal-Wallis and Dunn’s tests).

Given the influence of DA on animal displacement, we asked whether exogenous supplementation of this compound could protect from the neurotoxicity caused by cisplatin. Strikingly, we noticed that 5 and 10 mM DA rescue cisplatin-altered path range to WT standards, being 10 mM the most effective dosage (**Fig. 5B and Movie S3**). Therefore, our data suggest that DA protects from cisplatin-induced neurotoxicity in *C. elegans*.

## DISCUSSION

Our previous findings revealed that the conserved IIS pathway is involved in cisplatin response by activating its main transcription factor DAF-16/FOXO (García-Rodríguez et al., 2018). The IIS pathway, through diet, regulates a broad variety of processes, such as stress resistance, innate immunity, and metabolic adaptation in animals (Singh and Aballay, 2009; Yang and Hung, 2009; Murphy and Hu, 2013). Although high-glucose diets *per se* cause a myriad of phenotypes (such as reproductive alterations and aging) and metabolic changes (such as lipid composition and fat accumulation) (Alcántar-Fernández et al., 2018; Mejía-Martínez et al., 2017), we observed a synergistic impact on animal development between cisplatin and moderate and high glucose levels (40 mM and 80 mM, respectively). Interestingly, we also detected a hormetic effect against cisplatin-induced toxicity at low glucose dosage (10 mM). Similar mild resistance to cisplatin could contribute to the worst outcome of diabetic cancer patients with poor glycemic control (Bergamino et al., 2019). Thus, metabolism influences cisplatin response and consequently could be modulated for more efficient chemotherapy. In fact, adaptive metabolic responses to oxidative agents, including a reduction of the core biological processes, have been reported in distinct animal models (Shore et al., 2012; Ventura et al., 2009).

Mitochondrial BH3-only CED-13 protein protects from mtROS (Yee et al, in 2014), and consistently, we demonstrated the protective role of CED-13 against cisplatin-induced toxicity (García-Rodríguez et al.2018). Our results suggest that *ced-13* also protects from the synergistic toxicity of cisplatin with glucose and paraquat, pointing towards excessive cellular oxidation as the main consequence of such synergy. The UPR^mt^ pathway activation under cisplatin exposure further sustains the impact of cisplatin in mtROS. Since imbalance of OXPHOS complexes is one of the leading causes of UPR^mt^ pathway activation (Choi et al., 2015), we studied the *C. elegans* cellular respiration and quantified its reduction in the presence of cisplatin, glucose, and paraquat. Interestingly, we noticed that cisplatin exposure *per se* and in combination with glucose and PQ negatively impacts cellular respiration at different levels.

Mitochondria plays a crucial role in cisplatin cytotoxicity. In clinics, mitochondrial content has been proposed as a biomarker for platinum-based therapy response and, in counterpart, as a target for cisplatin resensitizing strategy (Cocetta et al., 2019). Mitochondrial accumulation of cisplatin triggers the imbalance of both mitochondrial redox and the signal crosstalk with the nucleus, altering cell metabolism (Choi et al., 2015). Moreover, mitochondrial dynamics are also relevant in cancer cells’ adaptation mechanism to stressful conditions, including chemotherapy (Cocetta et al., 2019). The link between cisplatin cytotoxicity and mitochondrial dynamics has also been described in yeasts (Inapurapu et al., 2017). Consistently, our findings indicate that mitochondria play a crucial role in cisplatin cytotoxicity and support the use of *C. elegans* as a model to investigate cellular and systemic responses to cisplatin in the context of a pluricellular organism.

Based on our previous published cisplatin transcriptomic signature in *C. elegans*, we hypothesized that catecholamine-*O*-transferases genes might be involved in cisplatin-induced neurotoxicity. By CRISPR-Cas9, we generated deletion alleles for those COMT family members with higher transcriptional activity at postembryonic stages. These null alleles are viable and suitable for functional studies of nematode behavior and locomotion. Additionally, we generated a new deletion allele for tyrosine hydroxylase *cat*-*2*. Thus, we built a toolkit with DA-related mutants to investigate the implication of DA in cisplatin neurotoxicity. After evaluating several DA-dependent effects in the distinct mutant backgrounds, we noticed that *comt*-*3*, −*4* and −*5* would present differential implications on DA catabolism. While *comt*-*5* has a major implication on body length, and *comt*-*4* and *comt*-*5* seem to affect animal locomotion, we did not find phenotypical alterations in the *comt*-*3* mutant. Functional redundancies among COMT genes in *C. elegans* should be explored in the future.

Recent publications have been described the relevance of catecholamine metabolism on animal behavior (Omura et al., 2012; Rodríguez-Ramos et al., 2017). Here, we focused on animal locomotion, specifically path range, to evaluate the neuroprotective effect of dopamine both in control and cisplatin conditions. First, we demonstrated that correct dopamine levels are essential to maintain normal levels of traveled distances, as evidenced in low- and high- DA mutants, *cat*-*2* and *comt*-*4* and −*5*, respectively, and when exogenous DA was administered. Interestingly, we found that exogenous DA also protects from cisplatin-induced neuronal damage. Similar findings have been reported in Zebrafish model, in which high DA levels (or alternatively L-miosine) not only protect from ototoxicity but also from nephrotoxicity induced by cisplatin but without affecting the toxicity in tumoral cells (Wertman et al., 2020). Although further experiments are needed to unravel the mechanisms beyond such protectiveness, there are two hypotheses to explain why increased DA levels mediate oto- and nephroprotection. On the one hand, DA binding to D1 to D5 receptors, present in the kidney and the mammalian inner ear, has been found to provide nephro- and cochlear nerve protection through increasing cAMP levels (Darrow et al., 2007; Gillies et al., 2015; Hans et al., 1990; Lendvai et al., 2011; Oestreicher et al., 1997; Ruel et al., 2001). Interestingly, D1, D2 and D3-like receptors are conserved in *C. elegans*. On the other hand, it has been proposed that dopamine could compete with cisplatin for organic cation transporters (OCT), particularly for hOCT2, which is highly expressed in the kidney and in the outer hair cells (Hucke and Ciarimboli, 2016). Indeed, treatment of patients with other cations, or disrupting of OCT2 in mice, ameliorate cisplatin-induced toxicities (Hucke et al., 2019; Meijer et al., 1982; Zhang and Sulzer, 2012). Our findings provide additional pieces of evidence for the potential utility of dopamine to mitigate cisplatin-induced neurotoxic effects avoiding the reduction of cisplatin doses.

## MATERIAL AND METHODS

### *Caenorhabditis elegans* strains and general methods

*C. elegans* strains were maintained using standard procedures (Stiernagle, 2006). Before conducting the experiments, animals were grown for at least two generations at the experimental temperature. Animals were synchronized using sodium hypochlorite (Porta-de-la-Riva et al., 2012). N2 was used as WT strain. Strains used in this study are listed in **Table S2**, pointing out the ones generated by CRISPR-Cas9 and the ones provided by the Caenorhabditis Genetics Center (CGC). Mutants generated by CRISPR-Cas9 were outcrossed twice, and all the used strains were genotyped before its use using MyTaq^TM^ DNA polymerase (Bioline) according to the manufacturer’s instructions. Primers used for genotyping are listed in **Table S3**.

### CRISPR-Cas9 (Nested CRISPR)

Guide RNAs were designed using both Benchling (www.benchling.com) and CCTop (Stemmer et al., 2015) online tools. All CRISPR-Cas9 mutant and reporter strains were obtained following a co-CRISPR strategy (Kim et al., 2014) using *dpy*-*10* as a marker to enrich for genome-editing events (Arribere et al., 2014). In the case of CER588 *cat*-*2(cer181[cat*-*2p::gfp::h2b 1*-*3])II*, Cas12a (Cpf1) were used instead Cas9. For the last, co-CRISPR was not viable beacause of the inneficient crRNA for *dpy*-*10*. In all the cases, mixes were injected into gonads of young adult P_0_ hermaphrodites using the XenoWorks Microinjection System and following standard *C. elegans* microinjection techniques. F_1_ progeny was screened by PCR using specific primers and F_2_ homozygotes were confirmed by Sanger sequencing. All the reagents for step 1 used in this study are listed in **Tables S4, S5**. The injection mix conditions for Nested CRISPR step 1 and 2 are described in (Vicencio et al., 2019) as well as universal sequences for step 2.

### Plates with special requirements

Plates with special requirements were prepared as follows: *(i)* Cisplatin plates: cisplatin (Accord) 1 mg/mL was used as a stock solution. For solid cisplatin plates preparation, 55 mm NGM plates, with 10 mL agar, were prepared. The next day, 600 μL cisplatin solution stock was added on the surface to reach the desire concentration. When dried, 300 μL overnight OP50 culture were seeded. *(ii)* High-glucose plates: D-(+)-Glucose powder (Sigma-Aldrich) was diluted in deionized water for stock solutions preparation at the desired concentration. 300 μL from the respective stock solution was added into the plates before been seeded and incubated overnight at room temperature. After incubation, plates were seeded with 300 μL overnight OP50 culture. *(iii)* Paraquat plates: Paraquat (Sigma-Aldrich) powder was resuspended in DMSO (Dimethyl sulfoxide) to reach 1M as a stock solution. Paraquat solution was added to NGM medium (still melted), mixed, and poured into 55 mm plates. 0.1 mM was used as final paraquat concentration. 300 μL overnight OP50 culture were seeded. (iv) Low peptone plates (tracking plates), for 1 L plates: 3 g sodium chloride, 20 g agar, 0.13 bactopeptone and 1L desionized water were mixed and autoclaved. Then, 3.5 cm plates where prepared with this solution plus standard concentration buffers (Stiernagle, 2006). Plates were seeded with a single drop in the middle of the surface from an overnight OP50 culture the day before the experiment.

### Body length assay

A synchronized population of L1-arrested larvae was cultured on NGM plates containing fresh OP50 and 60 μg/mL of cisplatin at 20°C. The body length of 50 animals for each condition was measured after 72 hours at the stereomicroscope using NIS-Elements 3.2 imaging system.

### Automated tracking of behavioral features

Synchronized animals were grown at 20°C in NGM plates from L1 for 34 hours. Then, animals were plated in 12-well plates (30 animals per well) containing liquid culture for each condition: control, cisplatin (Accord,1 mg/mL), or dopamine (Sigma-Aldrich) at the desired concentration for 24-hours treatment. 25 mL overnight OP50 culture diluted in 5 mL M9 were used as a source of food. Three biological replicates were prepared per each condition. The day of the experiment, animals were recovered and washed in M9 and finally seeded on tracking plates and let them for habituation for 10 minutes. Animals were recorded for 15 minutes using Tierpsy Tracker and data were extracted and analyzed using Tierpsy Tracker 2.0 software (Javer et al., 2018a, 2018b).

### Oxygen consumption rate (OCR) assessment

*C. elegans* respirometry profile was determined by measuring oxygen consumption rate (OCR) by Seahorse XFe96 Analyzer (Agilent). Optimized procedure proposed in Koopman et al. 2016 was followed with minor modifications. Oxygen consumption rate was calculated in N2 treated with 5 different conditions: 60 μg/mL cisplatin, 0.1 mM paraquat, 40 μM glucose, paraquat + cisplatin, glucose + cisplatin and H2O as a vehicle. The protocol described in Koopman et al., 2001 was followed with minor modifications. Plates with special requirements were prepare freshly as indicated above. We ensure that animals were well-synchronized and that the day of the experiment do not exceed the L3 stage since, after the L3/L4 molt, substantial differences in mitochondrial load would exist between younger and older animals affecting respirometry (Bratic et al., 2009; Tsang and Lemire, 2002).

### Graph plotting and statistical analysis

Data were plotted and differences between groups were analyzed using GraphPad Prism 8.0.

