## Supplementary tables and figures for "Insights into cisplatin-induced neurotoxicity and mitochondrial dysfunction in *Caenorhabditis elegans*"

#### Supplementary table 1: BLASTP analysis of *C. elegans* COMT proteins.

| Protein | Query cover (%) | Identity (%) |
|---|---|---|
| COMT-4 | 77 | 36.9 |
| COMT-3 | 17 | 74.47 |
| COMT-2 | 17 | 76.60 |
| COMT-5 | 17 | 65.96 |
| COMT-1 | 63 | 22.45 |

### Supplementary table 2: Strains used in this study.

| Strain | Genotype | Reference |
|---|---|---|
| CER494 | *comt-5(cer126 [comt-5p::gfp::h2b 1-3]) V* | This study |
| CER496 | *comt-4(cer128[comt-4p::gfp::h2b 1-3]) V* | This study |
| CER498 | *comt-3 (cer130[comt-3p::gfp::h2b 1-3) V* | This study |
| CER587 | *comt-3 (cer166[comt-3p::gfp::h2b 1-3); comt-4(cer128 [comt-4p::gfp::h2b 1-3]; comt-5(cer167 [comt-5p::gfp::h2b 1-3]) V* | This study |
| CER497 | *comt-4(cer128 [comt-4p::gfp::h2b 1-3] comt-3 (cer166[comt-3p::gfp::h2b 1-3) V* | This study |
| CER554 | *comt-4(cer157[comt-4p::GFP::H2B]) V* | This study |
| CER588 | *cat-2 (cer181 [cat-2p::gfp::h2b 1-3]) II* | This study |
| MT15620 | *cat-2 (n4547) II* | CGC |
| FX536 | *ced-13(tm536) X* | CGC |
| MD792 | *ced-13(sv32)* | CGC |

**Supplementary table 3: List of primers used for genotyping.**

| Gene | Allele | | Primer Fwd | Primer Rev |
|---|---|---|---|---|
| *comt-4* | *cer128*<br>*cer157* | | TCCAAAGTTCAGTTCGGAAG | CCAGAAATCGGACTTGATTGA |
| *comt-3* | *cer130*<br>*cer166* | | cttgttctctcggactatctgatag | TGTCTACTTTGCCCCCAATG |
| *comt-5* | *cer126*<br>*cer167* | | ccctccaaacagctattgaaacg | ACATGAGTTCCATCGCCAAGA |
| *cat-2* | *n4547*<br>*cer181* | WT | ctatgtgaagtcacacctgtc | gagatcacggatcacaagag |
| | *cer181* | Mut | | cttgctggaagtgtacttggtg |
| *ced-13* | *tm536* | | GTCAGGTGGCCCACGAAAC | CAGTACGTGCTTGATGCAC |
| | *sv32* | | | GGCAGTTGCTGAGACGTTG |

### Supplementary table 4: List of crRNAs used for Nested CRISPR step 1.

| Name | Generated allele | Sequence |
|---|---|---|
| *comt-4 5'* | *cer128* | TATTGTTGCCAAGAGTTACG |
| *comt-4 3'* | | TTCACTTCTTAAAAGCCATG |
| *comt-3 5'* | *cer130, cer160* | CGCAAAAAGCTACAAGAGCT |
| *comt-3 3'* | | TCGCTTTTAAGAAGTGAATT |
| *comt-5 5'* | *cer126, cer167* | TAAGGATGCCGATCCAGTGG |
| *comt-5 3'* | | AATTTCCAGAGCCTTCGCGG |
| *cat-2 5'* | *cer181* | CGTGATCCTCTCCAGAGCCC |
| *cat-2 3'* | | ACATTGTAATCGATATTTTC |

### Supplementary table 5: List of ssODN used for Nested CRISPR step 1.

| Locus | Generated allele | Sequence |
|---|---|---|
| *comt-4* | *cer128* | TTTCAGTTTTTTTTTCCGAAAAAAAAAAATGTCCAACccaagtttgtacaaaaaagcaggctccatgagtaaaggagaagaacttttcactggagagggaaccaaggccgtcaccaagtacacttccagcaagtaaATTAGGGGCTTTTTTTTTAATTTTGAATTATATTTA |
| *comt-3* | *cer130*, *cer166* | tttcagtttttttttccgaaaaaaaaaaATGTCCAACccaagtttgtacaaaaaagcaggctccatgagtaaaggagaagaacttttcactggagagggaaccaaggccgtcaccaagtacacttccagcaagtaaattaggggctttttttttaattttgaattatattta |
| *comt-5* | *cer126*, *cer167* | GTTGTCGCTAAGAGTTATCATAAGGATGCCGATCCAccaagtttgtacaaaaaagcaggctccatgagtaaaggagaagaacttttcactggagagggaaccaaggccgtcaccaagtacacttccagcaagtaaCGGTGGCTCCGTAGCTGACGAGAAAAGACGAGAAGA |
| *cat-2* | *cer181* | TCGTTGTTGGCGTGGGGACCCCTTGAAAAATTGGAAGAGGAAATGTTTTTGCGGTATATGCGGAGGCGGGGCagtaaaggagaagaacttttcactggagttgtcccaattgccgtgtctgagggaaccaaggccgtcaccaagtacacttccagcaagTGAAACCTAATTTACCTAATACTTTGCTAAACTAT |

**SUPPLEMENTARY FIGURES**

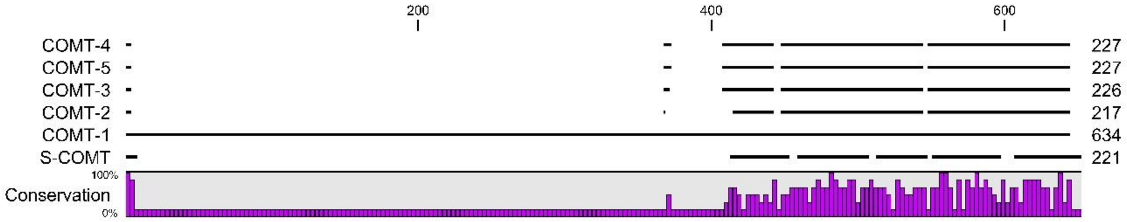


**Supplementary figure 1: Phylogenetic analysis of *C. elegans* COMT members. A.** Scheme of COMT-1 to -5 and S-COMT protein sequences (black lines) and conserved residues (purple bars). Alignments were illustrated with CLC Sequence Viewer 8.0.

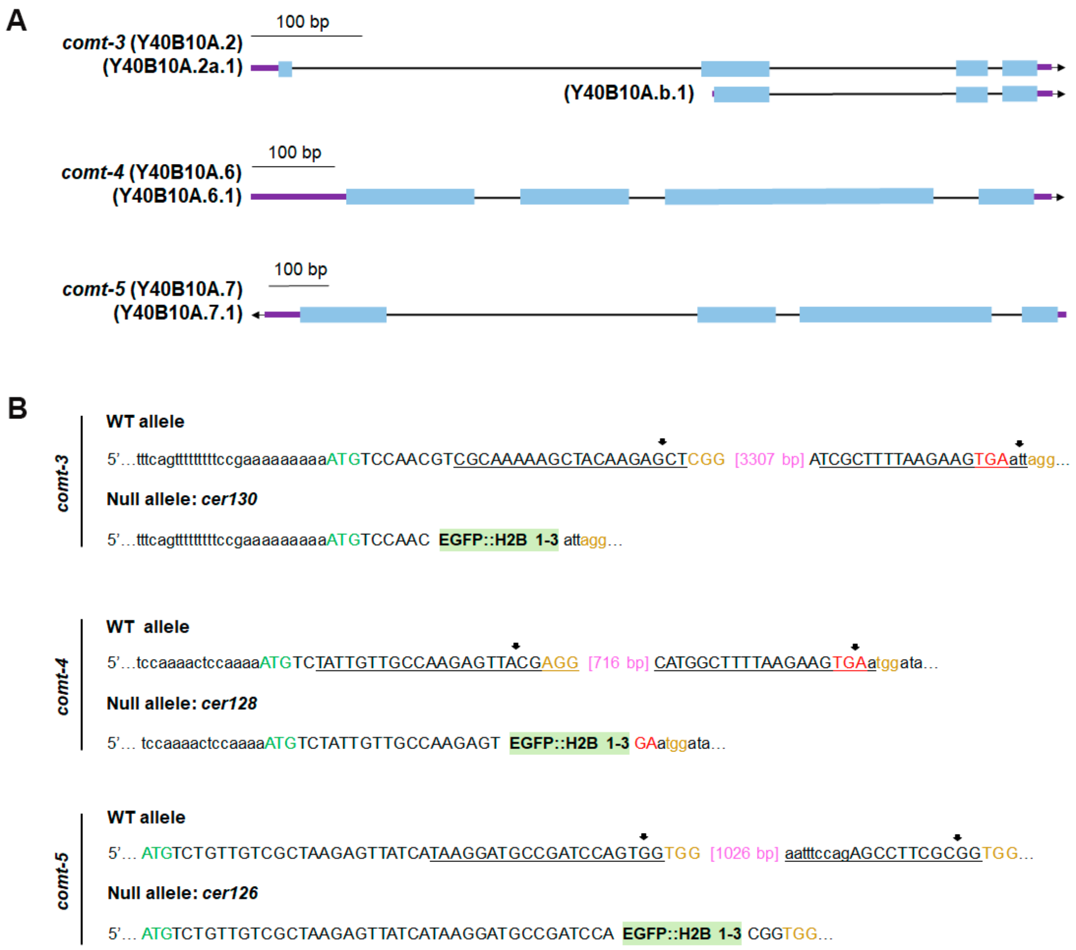


**Supplementary figure 2: *comt-3*, *comt-4* and *comt-5* loci and null allele molecular designs. A.** Schematic representation of *comt-3*, *comt-4* and *comt-5* transcripts. Introns are shown by black lines, 5´and 3´UTRs (untranslated regions) by purple bars and exons by light blue bars. **B.** Illustration of the sense strand sequences before and after homology-directed repair by CRISPR-Cas9 producing *cer130, cer128* and *cer126* null alleles. Start and stop codons are represented in green and red letters, respectively. PAM sequences are shown in yellow. 3´and 5´crRNA sequences are underlined and cut sites are shown by black arrows. The resulting deletion lengths are represented in pink and EGFP::H2B fragments 1 and 3 are represented by green shadow. Partial 5' and 3' UTR sequences are indicated in lowercase.

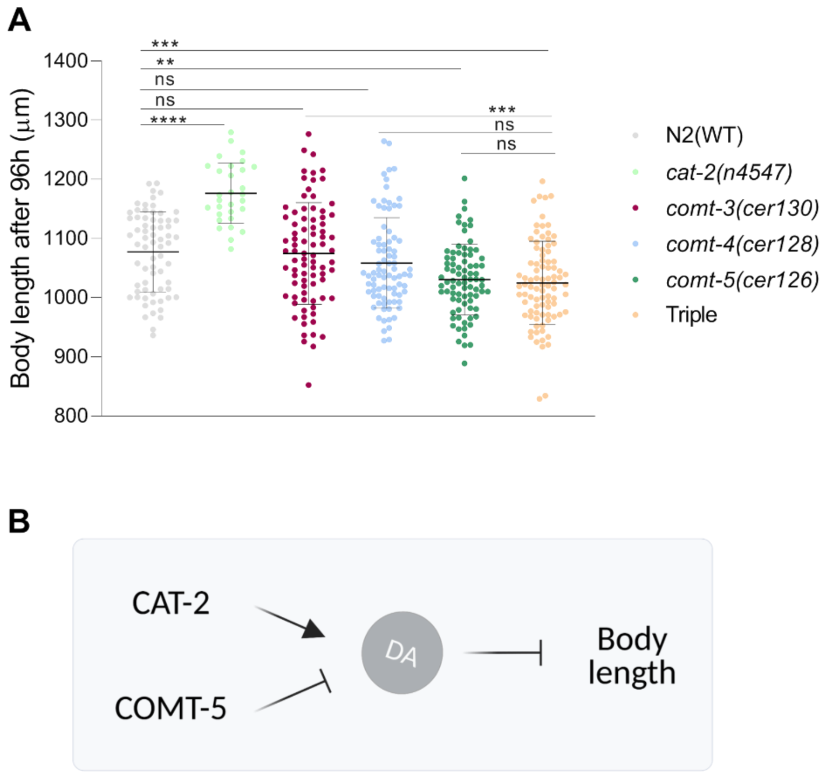


**Supplementary figure 3: Body length characterization of the dopamine signaling-related mutants.** Dots represent measurement of individual animals from three independent experiments, and black lines depict median and interquartile range. Strains used in these experiments are simple deletion alleles for *comt-3*, *comt-4*, *comt-5*, triple mutant for those, and deletion allele for *cat-2*. Statistical significances was assessed by one-way ANOVA (Krustal-Wallis and Dunn's tests). **, ***, **** and ns mean p<0.01, p<0.001, p<0.0001 and no significant, respectively.

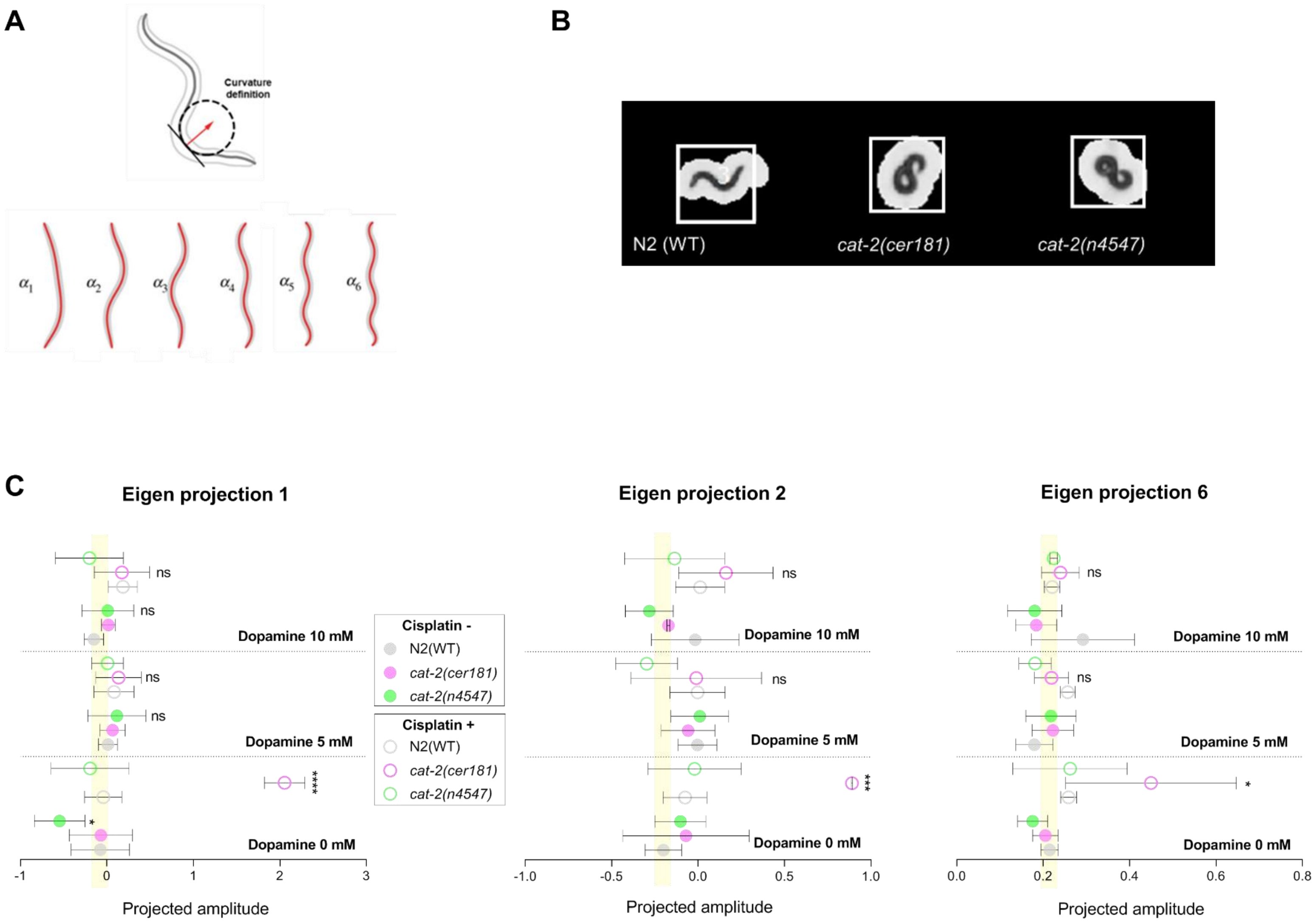
A
Curvature definition
α1
α2
α3
α4
α5
α6
B
N2 (WT)
cat-2(cer181)
cat-2(n4547)
C
Eigen projection 1
Eigen projection 2
Eigen projection 6
ns
Dopamine 10 mM
Dopamine 5 mM
Dopamine 0 mM
****
***
*
Cisplatin -
N2(WT)
cat-2(cer181)
cat-2(n4547)
Cisplatin +
N2(WT)
cat-2(cer181)
cat-2(n4547)
-1
0
1
2
3
-1.0
-0.5
0.0
0.5
1.0
0.0
0.2
0.4
0.6
0.8
Projected amplitude

**Supplementary figure 4: Body posture alteration of dopamine defective mutants induced by cisplatin. A.** On top of the figure, a worm diagram is illustrated, crossed by a line describing the shape. Curvature definition is given by the arc length function, represented by the red arrow. On the bottom, schematic representation of six eigenprojections accounting for almost the entire variance in body shape (Modified from Javer et al., 2018b). **B.** Representative images of animals exposed to cisplatin visualized by Tierpsy Tracker 2.0. **C.** Graph show projected amplitudes $\alpha_1$, $\alpha_2$ and $\alpha_6$ of dopamine-defective mutants. Circles represent the mean of two independent experiments and bars represent standard deviation. Yellow shadows point out the projected amplitude for wild-type animals in the absence of cisplatin nor dopamine. *, **, ns mean $p<0.1$, $p<0.01$ and not significant, respectively. Statistics were analyzed by one-way ANOVA (Krustal-Wallis and Dunn's tests).
